# Label-free characterisation of amyloids and alpha-Synuclein polymorphs by exploiting their intrinsic fluorescence property

**DOI:** 10.1101/2021.11.30.470691

**Authors:** Chyi Wei Chung, Amberley D. Stephens, Edward Ward, Yuqing Feng, Molly Jo Davis, Clemens F. Kaminski, Gabriele S. Kaminski Schierle

**Affiliations:** Department of Chemical Engineering and Biotechnology, Phillipa Fawcett Drive, University of Cambridge, CB3 0AS

**Keywords:** Fluorescence lifetime, intrinsic fluorescence, two-photon, 2P-FLIM, amyloid, fibril polymorphs

## Abstract

Conventional *in vitro* aggregation assays often involve tagging with extrinsic fluorophores which can interfere with aggregation. We propose the use of intrinsic amyloid fluorescence lifetime probed using two-photon excitation and represented by model-free phasor plots, as a label-free assay to characterise amyloid structure. Intrinsic amyloid fluorescence arises from structured packing of β-sheets in amyloids and is independent of aromatic-based fluorescence. We show that different amyloids (i.e., α-Synuclein (αS), β-Lactoglobulin and TasA) and different polymorphic populations of αS (induced by aggregation in salt-free and salt buffers mimicking the intra-/extracellular environments) can be differentiated by their unique fluorescence lifetimes. Moreover, we observe that disaggregation of pre-formed fibrils of αS and βLG leads to increased fluorescence lifetimes, distinct to those of their fibrillar counterpart. Our assay presents a medium-throughput method for rapid classification of amyloids and their polymorphs (the latter of which recent studies have shown lead to different disease pathology), and for testing small molecule inhibitory compounds.

## INTRODUCTION

Amyloid proteins aggregates share common characteristics, including a fibrillar morphology and cross-β sheet structure.^1^ The majority of *in vitro* studies on the kinetics of amyloid aggregation are fluorescence-based using extrinsic fluorophores with a fluorescence intensity readout, yet this presents issues when investigating small molecule inhibitors or fibril polymorphs (i.e., fibrils of different structures within the same amyloid species). Initially developed for histological stains of amyloid plaques in autopsies, common extrinsic fluorophores include Congo Red (CR) and Thioflavin T (ThT) which bind by intercalating between β-sheets of the amyloid of interest.^2^ Upon imaging in polarised light, CR-stained amyloids are revealed by apple-green birefringence; whereas increases in fluorescence intensity and quantum yield of ThT are observed with aggregation. However, the binding of both CR and ThT are affected by pH and ionic concentrations, ^3,4^ which must be strictly controlled under laboratory conditions. ThT based fluorescence assays can be affected by the binding of small inhibitory molecules; hence resulting in interference of fluorescence readings, due to either quenching effects between the molecule and ThT, or competitive binding to active sites on the amyloid protein.^5^ Furthermore, the presence of different disease-associated mutants of the Parkinson’s disease-related protein α-Synuclein (αS) leads to different ThT binding sites in its fibril polymorphs, and subsequently differences in ThT fluorescence intensity.^6^ Tagging recombinant proteins with fluorescent proteins or small dye-labels are also popular methods to study protein aggregation. Yet, the fluorescent protein tag can interfere with the excitation and emission of the ThT fluorescence^7^ and large fluorescent proteins can disrupt intramolecular bonding, sterically hinder interactions, and hence alter aggregation rates.^8^ As shown in a recent publication, even the presence of small dye molecules can influence the monomer incorporation into growing amyloid fibrils, thereby yielding polymorphic structures.^9^

Hence, there is motivation for the characterisation of amyloid protein fibrils in a label-free manner, which can be used to investigate potential inhibitors of amyloid aggregation and structural changes to the amyloids. In our previous work, we reported the phenomena of intrinsic amyloid fluorescence,^5,10–12^ as corroborated by similar studies from others.^13,14^ Characteristically, amyloid fibrils absorb light at wavelengths in the near UV range between 340 – 380 nm and emit fluorescence in the visible range between 400 – 450 nm. This phenomenon is believed to be caused by electron delocalisation due to the rich hydrogen bonding networks between and within the layers of β-sheets comprising an amyloid protein, along with the presence of short hydrogen bonds, resulting in the visible range emission upon UV excitation.^10,11^ It is noted that this phenomena is independent of intrinsic aromatic fluorescence as observed with aromatic amino acids (e.g., tyrosine and tryptophan) which exhibit both excitation and emission in the 260 – 280 nm UV range. Amyloid fibrils devoid of aromatic residues still show intrinsic non-aromatic fluorescence in the visible range upon UV excitation.^12,14^

Here, we explore the use of intrinsic amyloid fluorescence lifetime as a potential read out for aggregation states. We choose fluorescence lifetime over fluorescence intensity, as the first is a ratiometric parameter that is independent of excitation intensity, laser scattering, sample concentration and thickness.^15^ Several amyloids associated with neurodegenerative diseases feature fluorescence lifetimes in the nanosecond range, with measurements that dispute whether these are mono- or complex exponential in nature.^12,16^ Optimal excitation of amyloids is around 350 – 370 nm,^13^ wavelengths at which the power density of pulsed supercontinuum sources are very low. The alternative use of two-photon (2P) excitation which involves the absorption of two photons at twice the wavelength but half the energy,^17^ has inherent advantages. In contrast to single photon excitation which occurs through a cone of light down to the focal spot within a sample, 2P excitation (and hence any incurred photodamage) is primarily localised to the focal spot.^17,18^ This allows for imaging without a pinhole and is more suited for dimmer samples (e.g., intrinsic amyloid fluorescence) as no photons would be rejected due to the lack of a pinhole. The low scattering property of 2P makes it suitable for deeper penetration into samples. The most common implementation involves a femtosecond titanium-sapphire (Ti:S) laser such as the one used in this work, thereby making the technique proposed more accessible for researchers as the setup is commonly available on existing 2P microscopes primarily used for deep tissue imaging.^19–22^ Hence, we perform time-correlated single photon counting (TCSPC-) fluorescence lifetime imaging microscopy (FLIM), using 2P excitation. Moreover, we represent 2P-FLIM data on phasor plots, a global analysis approach that is efficient and parameter-free. ^23–25^ This involves the conversion time-domain TCSPC data into the frequency domain *via* a Fourier transform, thereby giving ‘phasors’ on a polar plot. This avoids pixel-by-pixel fitting of exponential decays (i.e., a requisite of conventional exponential fitting methods), therefore is highly efficiently and less computationally expensive. Moreover, mono- and complex exponential decay lifetimes can easily be distinguished based on their positions on the phasor plot.

To determine whether 2P-FLIM can differentiate between different amyloids, we used three model amyloids, e.g., β-lactoglobulin (βLG, a globular whey protein) TasA (a functional bacterial amyloid from *Bacillus subtilis*) and αS (the aggregation of which is a hallmark of Parkinson’s disease). We observe that they each have unique intrinsic fluorescence lifetimes, which can be used to distinguish between them. We validate our novel fluorescence lifetime measurements using circular dichroism (CD, which permits the analysis of protein secondary structure) and atomic force microscopy (AFM, which permits the characterisation of individual fibrils). In the amyloid field, the discovery of different fibril strain polymorphs is associated to different toxicity to cells,^26^ and potentially different disease outcomes. Hence, in order to better elucidate the pathology of amyloid misfolding diseases, it is useful to efficiently identify different fibril polymorphs.^27,28^ Currently, the best method to distinguish between these is cryogenic electron microscopy (cryoEM) due to its high resolution, yet it is a technique that not all researchers have access to and is expensive and low throughput. In the case of αS, we show different distributions of fibril polymorphs are formed in ‘no salt’ and ‘salt’ (i.e., mimicking the physiological environment in cells) conditions, which can be distinguished using 2P-FLIM measurements. We further show an increase fluorescence lifetime when the amyloid proteins are disaggregated, indicating structural changes to the amyloids leads to changes in fluorescence lifetime that can be tracked. We therefore provide a cheaper and higher-throughput technique to identify different amyloid fibrils.

## RESULTS

We initially performed structural and optical characterisation of the three fibrillar proteins of interest (**Figure 1**). Firstly, we used CD to determine the secondary structure of proteins. The method measures changes in the ellipticity of circularly polarised light when absorbed by different secondary structures (e.g., β-sheet, α-helix) of the protein. We observe that αS (pink) has the highest proportion of β-sheets (i.e., lowest mean residue ellipticity ∼ 220 nm) compared to both βLG (blue) and TasA (magenta) (**Figure 1a**). Monomeric αS is intrinsically disordered, but it undergoes structural alteration to β-sheets upon fibrillisation.^29^ On the other hand, βLG and TasA both contain β-sheets and α-helices in their monomeric form; both proteins have a decrease in α-helices and an increase in inter-molecular β-sheets upon aggregation.^30,31^ In order to perform physical characterisation on single fibrils we used AFM to analyse fibril morphology (**Figure 1b** and **Supplementary Figure 2**). From comparing height profiles, TasA fibrils are evidently shorter in height at 1.0±0.3 nm and without distinctive pitches, in comparison to βLG and αS, with average height profiles of 9.5±3.6 nm and 9.0±3.2 nm, respectively. For βLG and αS, there is a relatively large spread in the height of fibrils formed, which indicates the heterogeneity that exists within the same species sample. Single photon spectrofluorometric measurements reveal that the intrinsic fluorescence for each different amyloid has different optimal excitation and emissions wavelengths in the near UV and visible range respectively (βLG – ex 360 nm, em 430 nm, TasA – ex 350 nm, em 435 nm, and αS – 380 nm, em 425 nm, **Figure 1c**) and this suggests that they can be excited by 2P.

**Figure 1.**
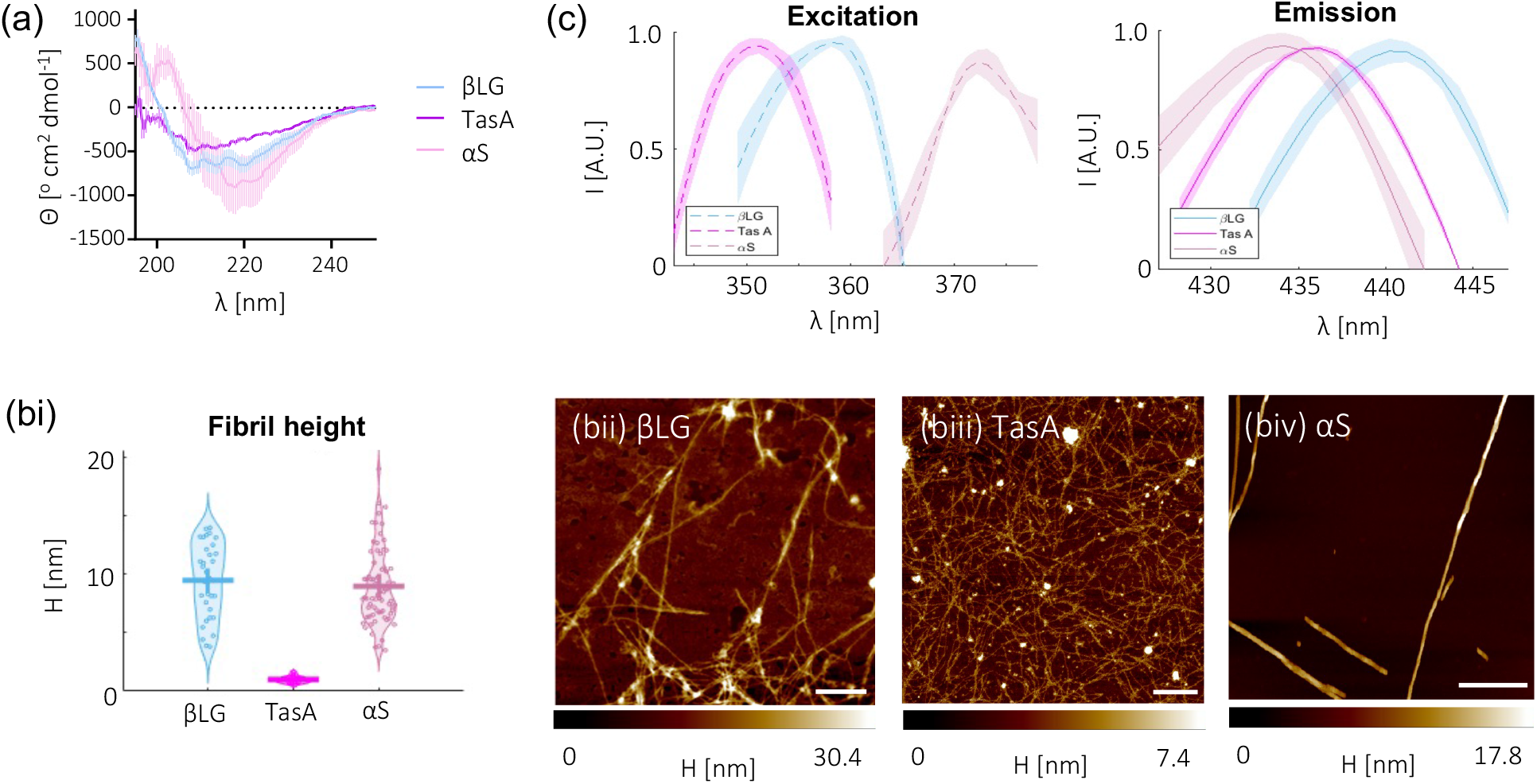
Different amyloid species can be differentiated by their spectral signatures. (a) CD spectra, displayed as mean ellipticity per residue (Θ), show that αS (pink) has a higher β- sheet content than βLG (blue) and TasA (magenta). Displayed are the average of 10 scans from three individual protein preparations. (b) Representative AFM images show the resulting amyloid fibrils have different morphologies. A height quantification is given in (bi). (bii) βLG fibrils in H_2_O are on average 9.5±3.6 nm in height (quoted as mean ± SD). (biii) TasA fibrils in 10 mM Tris pH 8 have no periodicity and are on average 1.0±0.3 nm in height. (biv) αS fibrils in 10 mM Tris pH 7.4 have mixed polymorphs with some fibrils displaying periodicity and others not. Their average height is 9.0±3.2 nm. The height profiles of a total of 33, 15 & 71 fibrils are analysed for βLG, TasA and αS respectively, based on 3 individual protein preparations. Additional AFM images are given in **Supplementary Figure 2**. (c) Excitation and emission spectra are measured between 340 to 400 nm with the emission set at peak emission, and emission spectra are measured between 400 to 460 nm with the excitation set at peak excitation for each protein. Excitation and emission peaks for each protein are, βLG – ex 360 nm, em 430 nm, TasA – ex 350 nm, em 435 nm, and αS – 380 nm, em 425 nm. Displayed are the average of three scans from three individual protein preparations.

We next investigated the intrinsic fluorescence lifetime signatures of the three amyloid fibril samples using 2P-FLIM. We also image the sample topography with AFM, as the diffraction limit on our 2P-FLIM system does not permit the visualisation of small fibrils. We deposit fibrils washed in dH_2_O at 100 μM onto clean glass coverslips which are then dried before imaging to provide a dense coverage of protein (**Figure 2a, AFM**). The lifetime of the intrinsic fluorescence emission reveal that all amyloids possess complex exponential with phasors that fall within the universal semicircle (i.e., which denotes mono-exponential lifetimes) of the phasor plot (**Figure 2b**) and in distinct positions from one another. Moreover, there are significant differences in their modulation (τ_M_) and phase (τ_ϕ_) lifetimes; for comparison of fluorescence lifetime, τ_M_ will be quoted henceforth as it is more sensitive than τ_ϕ_. (For an introduction to phasor plot analysis, see **Supplementary pp9—10**). We measure that βLG has the highest fluorescence lifetime at 1.7±0.2 ns, in comparison to TasA (0.96±0.02 ns) and αS (1.1±0.1 ns) (**Figure 2c**). We note that monomeric fluorescence is too weak to be detected on our 2P-FLIM system.

**Figure 2.**
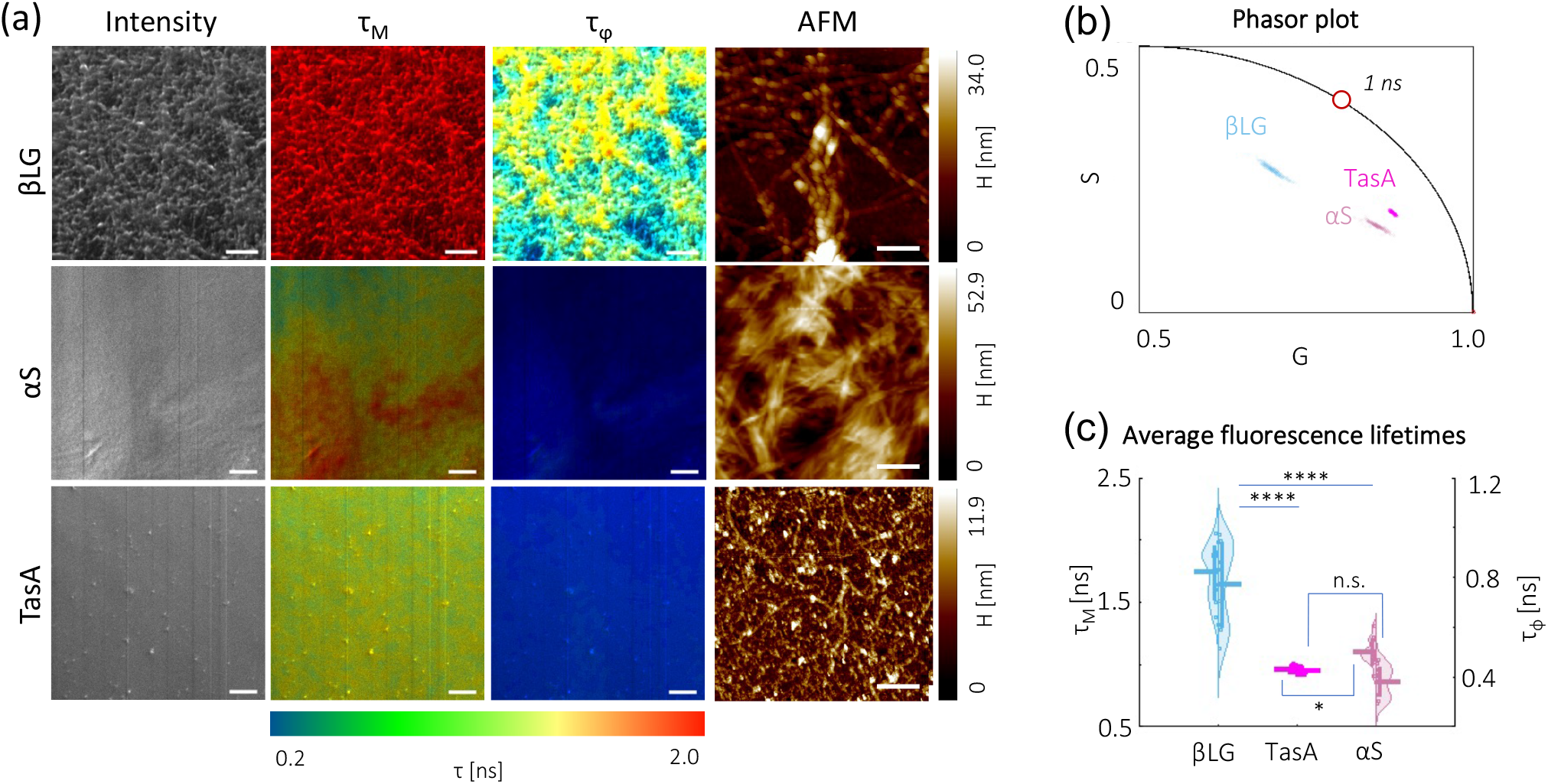
βLG, TasA and αS display different intrinsic fluorescent lifetime signatures. (a) Fluorescence intensity, fluorescence lifetime (i.e., both modulation τ_M_ and phase τ_ϕ_) and AFM representative images are shown. Their fluorescence lifetimes follow a multi-exponential decay, as seen from the differences in calculated τ_M_ and τ_ϕ_ (denoting that these phasors lie within the universal semicircle). Scale bars, 10 μm (FLIM) and 400 nm (AFM). (b,c) Phasor plots and average calculated fluorescence lifetimes which show that each amyloid has a distinctive lifetime, but that of βLG is significantly higher. Analysis is based on 10-12 images taken over 3 individual protein preparations. One-way ANOVA test (Holm-Sidak’s multiple comparison) where n.s. is not significant, * is p<0.05, and **** is p<0.0001.

It has been suggested that structurally different αS fibril polymorphs can lead to different synucleopathies, due to differences in membrane binding, seeding behaviour and toxicity.^32–34^ Results in **Figure 2** clearly show that different amyloids can be distinguished by their fluorescence lifetime signatures. Hence, this encouraged us to investigate if fluorescence lifetime is also responsive to more subtle structural changes, e.g., polymorphic variants that emerge for the same protein when aggregated under different buffer conditions. It has previously been shown that ‘no salt’ and ‘salt’ aggregation buffer conditions induce the formation of mixed populations of αS polymorphs, where fibrils formed in high salt conditions have distinct periodic pitches instead of flat ribbon structures.^35^ Our ‘no salt’ condition contains 10 mM Tris pH 7.4 (denoted as Tris) and two ‘salt’ conditions features the addition of 2 mM CaCl_2_ and 140 mM NaCl (CaCl_2_/NaCl, i.e., extracellular-mimicking) and 140 mM KCl (KCl, i.e., intracellular-mimicking). 2P-FLIM measurements show lowered fluorescence lifetimes for αS fibrils formed in KCl (0.95±0.09 ns) or CaCl_2_/NaCl salts (0.96±0.05 ns) compared to αS when aggregated in just Tris buffer (1.1±0.1 ns) (**Figure 3**). Although the magnitude of difference is slight compared to those between different amyloid species (**Figure 2**), this is as expected as there are less structural and molecular packing differences between the αS samples than between differing proteins. Moreover, their fluorescence spectra (**Supplementary Figure 3**) also show similar optimal excitation and emission wavelength.

**Figure 3.**
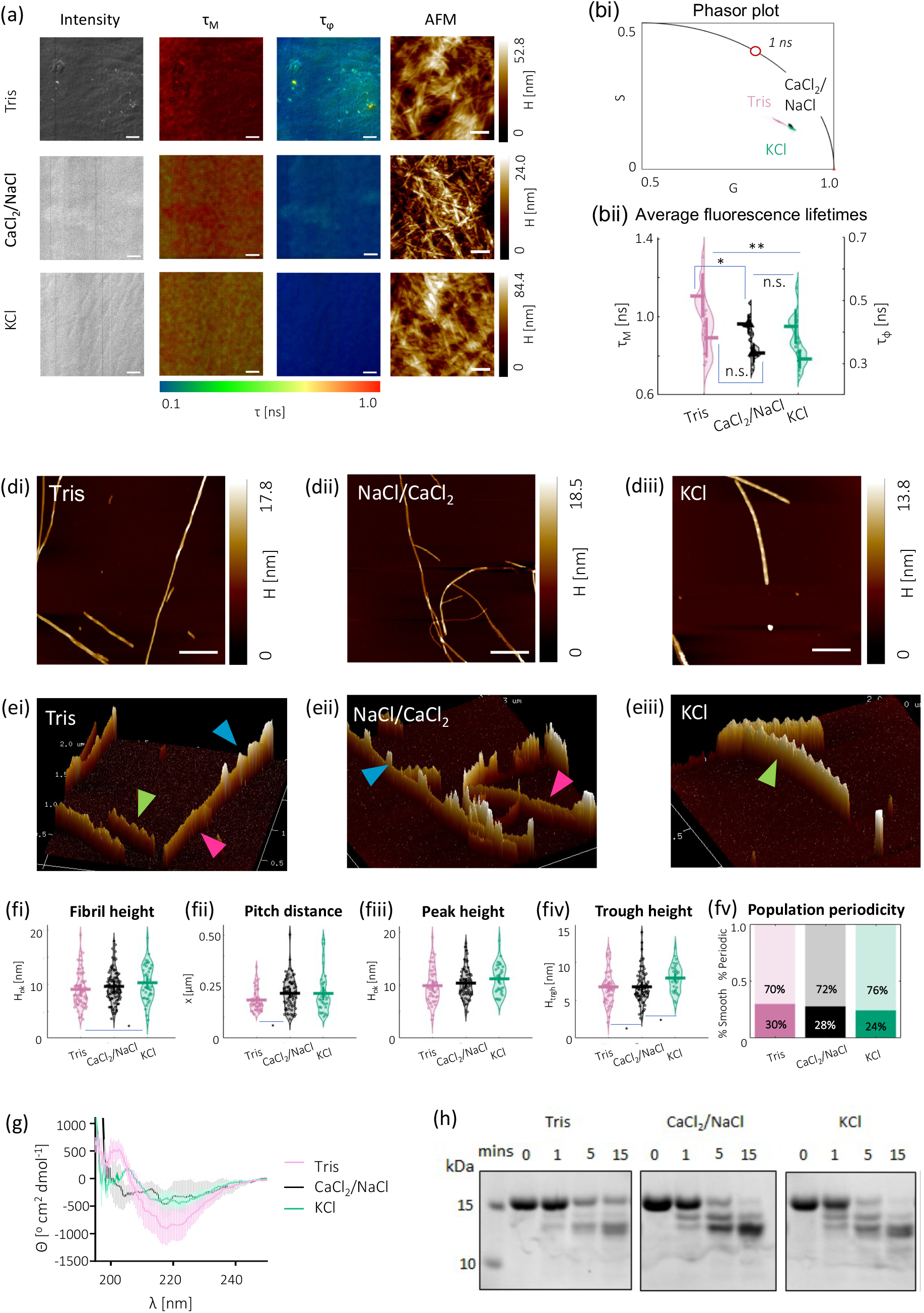
αS aggregated in salt buffers exhibit lower fluorescence lifetimes due to variation in the distribution of their polymorphs. (a) Fluorescence intensity, fluorescence lifetime (i.e., both modulation τ_M_ and phase τ_ϕ_) and AFM representatives are shown for αS fibrils formed in 20 mM Tris pH 7.4 (Tris), with 1 mM CaCl_2_ and 140 mM NaCl (CaCl_2_/NaCl), and with 140 mM KCl (KCl). Scale bars, 10 μm (FLIM) and 400 nm (AFM). (b,c) Phasor plots and average calculated fluorescence lifetimes which show that αS fibrils formed in salts have a lower average fluorescence lifetime compared to those formed in Tris only buffer. Analysis is based on 12 images taken over 3 individual protein preparations. One-way ANOVA test (Holm-Sidak’s multiple comparison) where n.s. is not significant, * is p<0.05, * is p<0.001, and **** is p<0.0001. (d) Shown are AFM images of individual fibrils in different buffers for morphological analysis. (e) 3D representations are shown to highlight structural differences. Different morphologies are indicated by coloured arrows, i.e., flat (pink), twisted periodic, from intertwined fibrils (blue) and periodic (green). Additional AFM images are given in **Supplementary Figure 4**. (f) Quantification of fibrils shows that there is great heterogeneity within each sample. In general, fibrils aggregated in salt buffers are higher with a lower frequency of pitches. The addition of salts slightly increases the chance of periodic fibrils over flat ones. The height profiles of a total of 71, 97 & 47 fibrils are analysed for αS aggregated in Tris, CaCl_2_/NaCl and KCl, respectively, based on 3 protein preparations. One-way ANOVA test (Holm-Sidak’s multiple comparison) where ns is not significant, * is p<0.05, * is p<0.001, and **** is p<0.0001. (g) CD of 2.5 μM of each protein fibrils show that αS in Tris has a higher β-sheet content compared to αS in 140 mM KCl and 140 mM NaCl and CaCl_2_. 10 scans of each sample were taken, and the average is shown for three protein sample preparations, with buffer only signals subtracted (h) 100 μM of αS in each buffer condition was incubated in proteinase K for 0, 1, 5 and 15 mins. Monomeric αS has a molecular weight of ∼14.4 kDa, degradation patterns of the three samples show a similar band profile, but the intensities differ, indicating differences in cleavage rates. A second repeat is shown in **Supplementary Figure 5**, with similar proteolysis profiles.

We then further characterise the three αS fibril samples to determine whether they are truly structurally and/or morphologically different. 2D and 3D AFM images show several different αS fibril polymorphs within each buffer condition (**Figure 3d-e** and **Supplementary Figure 4**), these polymorphs can be classified as either smooth (pink arrows), periodic (green arrows) or twisted periodic, likely arising from two fibrils twisting around each other (blue arrows). We performed single fibril analysis based on AFM images, to classify the height distribution and the prevalence of periodicity within each sample. As before, we observe a wide range of heights from fibrils within the same sample, with an average height of 9.0±3.2 nm (Tris), 9.6±2.9 nm (CaCl_2_/NaCl), and 10.2±3.3 nm (KCl) (**Figure 3fi**). The addition of salts promotes the formation of higher and intertwined fibrils, of which there is a greater proportion of those being periodic (i.e., 72% & 76% for CaCl_2_/NaCl and KCl, respectively in comparison to 70% for Tris, **Figure 3fv**). This is most apparent in the αS fibrils formed in KCl, where the fibril height distribution is more bimodal, showing single fibril height and double fibril height (**Figure 3fi**).

CD measurements show that αS aggregated in salt buffers have a decreased β-sheet content compared to those in Tris (**Figure 3g**). Furthermore, differences in fibril proteolysis profiles can be used to indicate a different fibril structure and core due to differences in accessibility of the protease.^36^ Limited proteolysis of the αS fibrils in the three buffers with proteinase K shows similar digestion profiles, but differing band intensity, indicating similarities in core structure, but differences in fibril packing and accessibility of proteinase K to the cleavage sites in the different fibril samples (**Figure 3h**, with a repeat shown in **Supplementary Figure 5**). Therefore, our sample characterisation supports structural differences in the fibrils formed under different salt conditions which we observe to possess different fluorescence lifetimes.

Lastly, to validate the use of intrinsic fluorescence lifetime as an *in vitro* label-free aggregation assay, we disaggregate fibrillar αS and βLG by mixing the samples with hexafluoro-2-propanol (HFIP), a solvent typically used to monomerise proteins before aggregation. We observe the formation of shortened fibrils and oligomers in αS and βLG, respectively, in AFM images (**Figure 4a**, AFM). Correspondingly, these disaggregated structures lead to significantly increased fluorescence lifetimes, especially in the case of oligomeric βLG (2.6±0.1 ns from 1.7±0.2 ns) and less so for αS (1.2±0.06 ns from 1.1±0.1 ns) (**Figure 4b-c**). As the intrinsic amyloid fluorescence could still be detected from both samples, this insinuates there is still β-sheet stacking present in the disaggregated structures of βLG and αS, yet a change in the stacking or arrangement during partial disaggregation has led to a change in intrinsic fluorescence lifetime.

**Figure 4.**
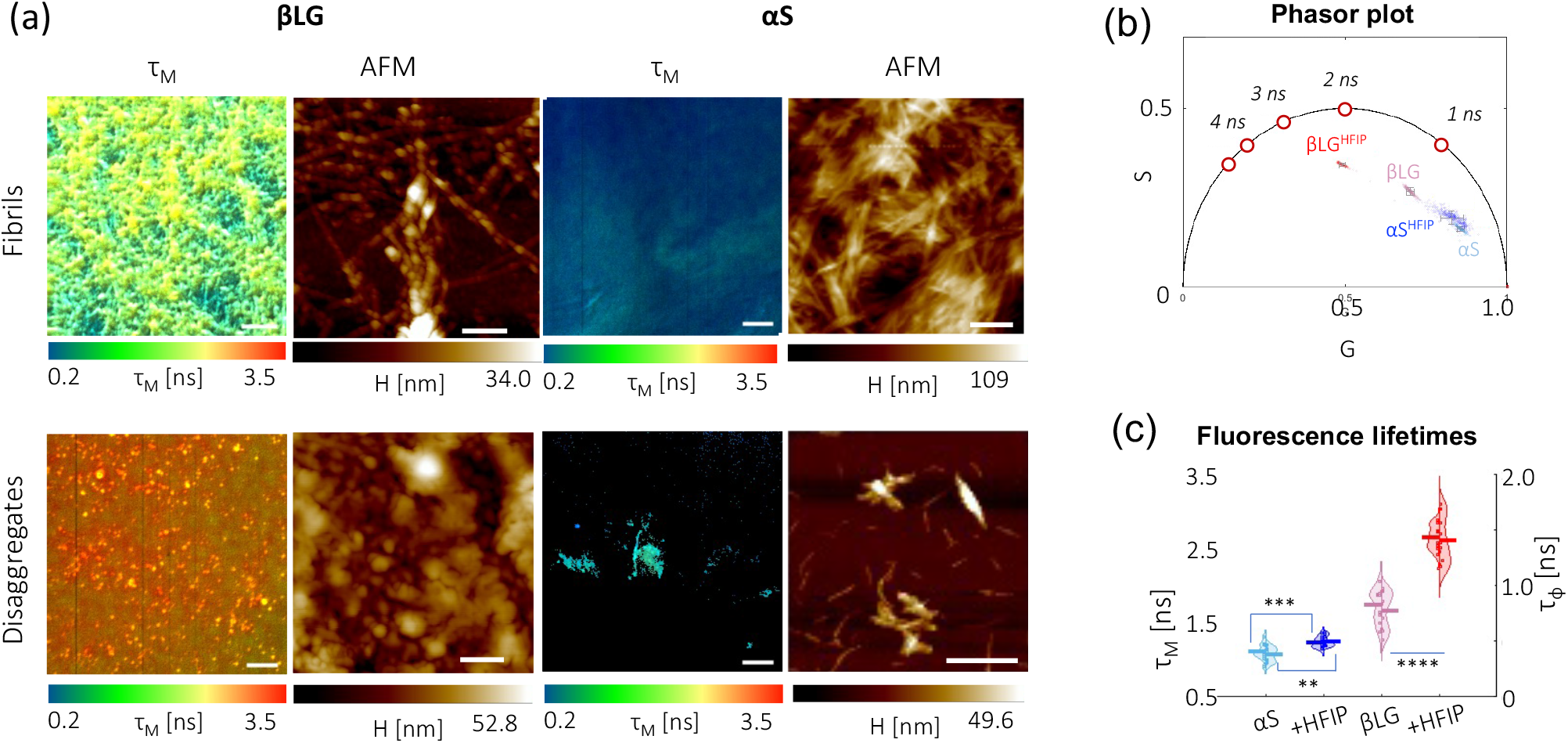
The fluorescence lifetime change of different amyloid proteins upon aggregation is reversible with disaggregated structures of αS and βLG having higher fluorescence lifetimes than their fibrillar counterparts. (a) Fluorescence lifetimes and AFM images comparing αS and βLG fibrils before and after disaggregation by HFIP. Scale bars, 10 μm (FLIM) and 400 nm (AFM). (b,c,) Phasor plot and average fluorescence lifetimes show a significant decrease in fluorescence lifetime after the addition of HFIP to disaggregate the fibrils for both αS and βLG. One-way ANOVA test (Holm-Sidak’s multiple comparison) where n.s. is not significant, * is p<0.05, and **** is p<0.0001.

## DISCUSSION

There is a need for label-free techniques to identify and monitor aggregation of amyloid proteins that is currently unmet. Here, we use a combination of structural and morphological techniques to validate the use of 2P-FLIM in identifying different amyloid protein fibrils, their polymorphs and their disaggregated states using fluorescence lifetime imaging. We investigate three amyloid proteins: βLG, a milk whey protein which is commonly used as a model amyloid protein, as it is inexpensive and readily available; TasA, a functional amyloid of *B. subtilis* involved in adherence and formation of the extracellular scaffold of biofilms; and αS, whose aggregation is a hallmark of synucleinopathies, such as Parkinson’s Disease. The fluorescence lifetime of conventional fluorophores, e.g., GFP, are influenced by the surrounding environment.^37,38^ The field of intrinsic non-aromatic fluorescence is comparatively young, therefore studies into differences in amyloid structures and interaction with the local environment have not yet been fully conducted yet may well be influenced by fibril packing and environmental interactions, which can lead to unique lifetimes for different proteins and polymorphs. Here, we efficiently represent the unique intrinsic fluorescence lifetime signatures of different amyloids, using model-free phasor plot analysis.

We then investigated whether 2P-FLIM could be used to identify fibril polymorphs of the same protein. The fibril formation of αS is implicated with several neuropathological diseases, including but not limited to, Parkinson’s disease, dementia with Lewy bodies, and multiple system atrophy.^39^ As aforementioned, cryoEM which boasts of a 3.7 Å resolution,^27,40^ is currently the gold standard technique for distinguishing and classifying amyloid (including αS) polymorphs, yet is an expensive technique that requires skilled users.^1^ Moreover, cryoEM has low throughput, hence it is unsuited for small molecule screening and cannot be used to investigate changes in protein states as they aggregate or change under different environmental conditions. Here, we use 2P-FLIM and show that αS fibrils formed in salt buffers mimicking the intracellular and extracellular have a slightly quenched fluorescence lifetime compared to the polymorphs formed without salt. From a structural perspective, noting that the fibrils are washed in H_2_O prior to 2P-FLIM imaging to remove salt ions, this quenching effect could be attributed to differences in the fibril packing (from limited proteolysis) or β-sheet structure (from CD) leading to the formation of different fibril polymorphs, that appear to have higher fibrils and less frequent pitches (from AFM). While we identify bigger differences between the ‘no salt’ Tris sample and the salt samples, there is a slight, but insignificant difference in the fluorescence lifetimes which is lower for αS fibrils formed in KCl than in CaCl_2_/NaCl. The two salt conditions were chosen to mimic intracellular and extracellular conditions, where in the latter calcium concentrations are higher. We, along with others, have previously shown that αS specifically binds calcium at its negatively charged C-terminus, this leads to altered monomeric structures and fibril polymorphs which may be important in cases of αS spreading from neuron to neuron.^41,42^ It is not known whether the structures formed *in vitro* are those that are formed *in vivo*, ^43^ in addition to whether different structures arise when aggregated in the intracellular and extracellular space. Further structural analysis using mutants and computational simulations may be able to pin-point the mechanisms that derive differences in fluorescence lifetime.

To quantify structural differences, we show that disaggregating pre-formed fibrils of αS and βLG result in an increase to their measured fluorescence lifetimes. We believe this stems from the looser packing and reduction β-sheets in within the disaggregated structures (i.e., smaller fibrils for αS and oligomers for βLG).^44^

## CONCLUSION

We validate that intrinsic amyloid fluorescence lifetime can be used as a label-free method to characterise different amyloid proteins, as well as the distribution within polymorphic populations of αS and disaggregated structures. Our current work comprises of observations on intrinsic amyloid fluorescence, which we find is affected by several different factors, e.g., β-sheet content and molecular packing. 2P-FLIM and efficient phasor plot analysis of fluorescence lifetimes maybe useful if applied to drug screening for amyloid protein targeting compounds, as 2P-FLIM can circumvent issues with small molecule interference with fluorescence intensity-based assays. To complement our findings, we believe that computational studies on the molecular structure of these amyloids at an atomistic scale that permit the study of electron transitions would be needed to establish causative links between structure and the unique fluorescence lifetime signatures amyloid possess. In general, intrinsic amyloid fluorescence lifetime in conjunction with fit-free phasor plot analysis provides a medium-throughput, efficient and label-free method to distinguish between different amyloids and their polymorphs.

## ASSOCIATED CONTENT

### Supporting Information

Supplementary materials (PDF) containing experimental methods and supplementary figures 1-5.

## AUTHOR INFORMATION

### Author Contributions

^‡^ C.W.C. and A.D.S contributed equally. G.S.K.S., C.W.C. and A.D.S conceived the manuscript. C.W.C, A.D.S, Y.F. and M.J.D. prepared protein for experiments. C.W.C. and E.W. built the 2P-FLIM, C.W.C collected 2P-FLIM data, C.W.C analysed 2P-FLIM data. C.W.C. collected excitation and emission spectra. A.D.S. collected CD data. C.W.C and A.D.S. collected AFM data, C.W.C analysed AFM data. A.D.S. performed limited proteolysis. C.F.K. provided equipment. All authors contributed to paper writing. All authors have given approval to the final version of the paper. All raw data is available on request and in the Cambridge University Repository (Apollo ID: A48E00D8-68DB-45B7-A368-406503E36D23). All analysis scripts are available on request.

## ACKNOWLEDGMENT

C.W.C. jointly funded by the Cambridge Trust and Wolfson College for her PhD. G.S.K.S. acknowledges funding from the Wellcome Trust (065807/Z/01/Z) (203249/Z/16/Z), the UK Medical Research Council (MRC) (MR/K02292X/1), Alzheimer Research UK (ARUK) (ARUK-PG013-14), Michael J Fox Foundation (16238) and Infinitus China Ltd. M.J.D. was funded by NanoDTC ESPSRC Grant EP/S022953/1. We thank Naunehal Matharu for preliminary CD data not in the final manuscript and Maria Zacharopoulou for discussions on αS fibril polymorphs.

## ABBREVIATIONS

2P: two-photon
αS: α-Synuclein
βLG: β-Lactoglobulin
τ_M_: modulation lifetime
τ_ϕ_: phase lifetime
AFM: atomic force microscopy
CD: circular dichroism
CR: Congo Red
cryoEM: cryogenic electron microscopy
FLIM: fluorescence lifetime imaging microscopy
ThT: Thioflavin T
Ti:S: titanium-sapphire
TCSPC: time-correlated single photon counting.

## REFERENCES

(1) Guerrero-Ferreira, R.,, Kovacik, L.,, Ni, D.,, Stahlberg, H. New Insights on the Structure of Alpha-Synuclein Fibrils Using Cryo-Electron Microscopy. Curr. Opin. Neurobiol. 2020, 61, 89–95. https://doi.org/10.1016/j.conb.2020.01.014.

(2) Biancalana, M.,, Koide, S. Molecular Mechanism of Thioflavin-T Binding to Amyloid Fibrils. Biochimica et Biophysica Acta - Proteins and Proteomics 2010, 1804 (7), 1405–1412. https://doi.org/10.1016/j.bbapap.2010.04.001.

(3) Mikalauskaite, K.,, Ziaunys, M.,, Sneideris, T.,, Smirnovas, V. Effect of Ionic Strength on Thioflavin-T Affinity to Amyloid Fibrils and Its Fluorescence Intensity. Int. J. Mol. Sci. 2020, 21 (23), 8916. https://doi.org/10.3390/ijms21238916.

(4) Hackl, E. v.,, Darkwah, J.,, Smith, G.,, Ermolina, I. Effect of Acidic and Basic PH on Thioflavin T Absorbance and Fluorescence. Eur. Biophys. J. 2015, 44 (4), 249–261. https://doi.org/10.1007/S00249-015-1019-8/TABLES/3.

(5) Pinotsi, D.,, Buell, A. K.,, Dobson, C.M.,, Kaminski Schierle, G. S.,, Kaminski, C. F. A Label-Free, Quantitative Assay of Amyloid Fibril Growth Based on Intrinsic Fluorescence. Chem. Bio. Chem. 2013, 14 (7), 846–850. https://doi.org/10.1002/cbic.201300103.

(6) Sidhu, A.,, Vaneyck, J.,, Blum, C.,, Segers-Nolten, I.,, Subramaniam, V. Polymorph-Specific Distribution of Binding Sites Determines Thioflavin-T Fluorescence Intensity in α-Synuclein Fibrils. Amyloid 2018, 25 (3), 189–196. https://doi.org/10.1080/13506129.2018.1517736.

(7) Anderson, V. L.,, Webb, W. W. Transmission Electron Microscopy Characterization of Fluorescently Labelled Amyloid β 1-40 and α-Synuclein Aggregates. BMC Biotechnol. 2011, 11 (1), 1–10. https://doi.org/10.1186/1472-6750-11-125/FIGURES/6.

(8) Afitska, K.,, Fucikova, A.,, Shvadchak, V. v.,, Yushchenko, D. A. Modification of C Terminus Provides New Insights into the Mechanism of α-Synuclein Aggregation. Biophys. J. 2017, 113 (10), 2182–2191. https://doi.org/10.1016/J.BPJ.2017.08.027.

(9) Cosentino, M.,, Canale, C.,, Bianchini, P.,, Diaspro, A. AFM-STED Correlative Nanoscopy Reveals a Dark Side in Fluorescence Microscopy Imaging. Sci. Adv. 2019, 5 (6), eaav8062. https://doi.org/10.1126/sciadv.aav8062.

(10) Pinotsi, D.,, Grisanti, L.,, Mahou, P.,, Gebauer, R.,, Kaminski, C. F.,, Hassanali, A.,, Kaminski Schierle, G. S. Proton Transfer and Structure-Specific Fluorescence in Hydrogen Bond-Rich Protein Structures. J. Am. Chem. Soc. 2016, 138 (9), 3046–3057. https://doi.org/10.1021/jacs.5b11012.

(11) Stephens, A. D.,, Qaisrani, M. N.,, Ruggiero, M. T.,, Mirón, G. D.,, Morzan, U. N.,, Lebrero, M. C.G.,, Jones, S. T. E.,, Poli, E.,, Bond, A. D.,, Woodhams, P. J.,, Kleist, E. M.,, Grisanti, L.,, Gebauer, R.,, Zeitler, J. A.,, Credgington, D.,, Hassanali, A.,, Schierle, G. S. K. Short Hydrogen Bonds Enhance Nonaromatic Protein-Related Fluorescence. Proc. Natl. Acad. Sci. U. S. A. 2021, 118 (21). https://doi.org/10.1073/PNAS.2020389118.

(12) Chan, F. T. S.,, Kaminski Schierle, G. S.,, Kumita, J. R.,, Bertoncini, C. W.,, Dobson, C. M.,, Kaminski, C. F. Protein Amyloids Develop an Intrinsic Fluorescence Signature during Aggregation. Analyst 2013, 138 (7), 2156–2162. https://doi.org/10.1039/c3an36798c.

(13) Shukla, A.,, Mukherjee, S.,, Sharma, S.,, Agrawal, V.,, Kishan, K. V. R.,, Guptasarma, P. A Novel UV Laser-Induced Visible Blue Radiation from Protein Crystals and Aggregates: Scattering Artifacts or Fluorescence Transitions of Peptide Electrons Delocalized through Hydrogen Bonding? Arch. Biochem. Biophys. 2004, 428, 144–153. https://doi.org/10.1016/j.abb.2004.05.007.

(14) del Mercato, L. L.,, Pompa, P. P.,, Maruccio, G.,, della Torre, A.,, Sabella, S.,, Tamburro, M.,, Cingolani, R.,, Rinaldi, R. Charge Transport and Intrinsic Fluorescence in Amyloid-like Fibrils. Proc. Natl. Acad. Sci. U. S. A. 2007, 104 (46), 18019–18024. https://doi.org/10.1073/pnas.0702843104.

(15) Becker, W. Fluorescence Lifetime Imaging - Techniques and Applications. J. Microsc. 2012, 247 (2), 119–136. https://doi.org/10.1111/j.1365-2818.2012.03618.x.

(16) Hecker, L.,, Wang, W.,, Mela, I.,, Fathi, S.,, Poudel, C.,, Soavi, G.,, Huang, Y. Y. S.,, Kaminski, C. F. Guided Assembly and Patterning of Intrinsically Fluorescent Amyloid Fibers with Long-Range Order. Nano Lett. 2021, 21 (2), 938–945. https://doi.org/10.1021/ACS.NANOLETT.0C03672.

(17) Denk, W.,, Strickler, J. H.,, Webb, W. W. Two-Photon Laser Scanning Fluorescence Microscopy. Science 1990, 248 (4951), 73–76. https://doi.org/10.1126/SCIENCE.2321027.

(18) Benninger, R. K. P.,, Piston, D. W. Two-Photon Excitation Microscopy for the Study of Living Cells and Tissues. Curr. Protoc. Cell Biol. 2013, 59 (1), 4.11.1-4.11.24. https://doi.org/10.1002/0471143030.CB0411S59.

(19) Larson, A. M. Multiphoton Microscopy. Nat. Photonics 2010, 5 (1), 1–1. https://doi.org/10.1038/nphoton.an.2010.2.

(20) Moulton, P. F. Spectroscopic and Laser Characteristics of Ti:Al2O3. J. Opt. Soc. Am. B 1986, 3 (1), 125–133. https://doi.org/10.1364/JOSAB.3.000125.

(21) Svoboda, K.,, Yasuda, R. Principles of Two-Photon Excitation Microscopy and Its Applications to Neuroscience. Neuron 2006, 50 (6), 823–839. https://doi.org/10.1016/J.NEURON.2006.05.019.

(22) Miller, D. R.,, Jarrett, J. W.,, Hassan, A. M.,, Dunn, A. K. Deep Tissue Imaging with Multiphoton Fluorescence Microscopy. Curr. Opin. Biomed. Eng. 2017, 4, 32–39. https://doi.org/10.1016/J.COBME.2017.09.004.

(23) Ranjit, S.,, Malacrida, L.,, Jameson, D. M.,, Gratton, E. Fit-Free Analysis of Fluorescence Lifetime Imaging Data Using the Phasor Approach. Nat. Protoc. 2018, 13 (9), 1979–2004. https://doi.org/10.1038/s41596-018-0026-5.

(24) Liao, S.-C.,, Sun, Y.,, Coskun, U. FLIM Analysis Using the Phasor Plots; Champaign, 2014.

(25) Digman, M. A.,, Caiolfa, V. R.,, Zamai, M.,, Gratton, E. The Phasor Approach to Fluorescence Lifetime Imaging Analysis. Biophys. J. 2008, 94 (2), L14. https://doi.org/10.1529/BIOPHYSJ.107.120154.

(26) Peng, C.,, Gathagan, R. J.,, Lee, V. M. Y. Distinct α-Synuclein Strains and Implications for Heterogeneity among α-Synucleinopathies. Neurobiol. Disease 2018, 109 (Pt B), 209–218. https://doi.org/10.1016/j.nbd.2017.07.018.

(27) Li, B.,, Ge, P.,, Murray, K. A.,, Sheth, P.,, Zhang, M.,, Nair, G.,, Sawaya, M. R.,, Shin, W. S.,, Boyer, D. R.,, Ye, S.,, Eisenberg, D. S.,, Zhou, Z. H.,, Jiang, L. Cryo-EM of Full-Length α-Synuclein Reveals Fibril Polymorphs with a Common Structural Kernel. Nat. Commun. 2018, 9 (1), 1–10. https://doi.org/10.1038/s41467-018-05971-2.

(28) Guerrero-Ferreira, R.,, Kovacik, L.,, Ni, D.,, Stahlberg, H. New Insights on the Structure of Alpha-Synuclein Fibrils Using Cryo-Electron Microscopy. Curr. Opin. Neurobiol. 2020, 61, 89–95. https://doi.org/10.1016/j.conb.2020.01.014.

(29) Giasson, B. I.,, Murray, I. V. J.,, Trojanowski, J. Q.,, Lee, V. M. Y. A Hydrophobic Stretch of 12 Amino Acid Residues in the Middle of α-Synuclein Is Essential for Filament Assembly. J. Biol. Chem. 2001, 276 (4), 2380–2386. https://doi.org/10.1074/JBC.M008919200.

(30) Ioannou, J. C.,, Donald, A. M.,, Tromp, R. H. Characterising the Secondary Structure Changes Occurring in High Density Systems of BLG Dissolved in Aqueous PH 3 Buffer. Food Hydrocolloids 2015, 46, 216–225. https://doi.org/10.1016/J.FOODHYD.2014.12.027.

(31) Chai, L.,, Romero, D.,, Kayatekin, C.,, Akabayov, B.,, Vlamakis, H.,, Losick, R.,, Kolter, R. Isolation, Characterization, and Aggregation of a Structured Bacterial Matrix Precursor. J. Biol. Chem. 2013, 288 (24), 17559. https://doi.org/10.1074/JBC.M113.453605.

(32) Landureau, M.,, Redeker, V.,, Bellande, T.,, Eyquem, S.,, Melki, R. The Differential Solvent Exposure of N-Terminal Residues Provides “Fingerprints” of Alpha-Synuclein Fibrillar Polymorphs. J. Biol. Chem. 2021, 296. https://doi.org/10.1016/J.JBC.2021.100737.

(33) Meade, R. M.,, Williams, R. J.,, Mason, J. M. A Series of Helical α-Synuclein Fibril Polymorphs Are Populated in the Presence of Lipid Vesicles. NPJ Parkinson’s Disease 2020, 6 (1). https://doi.org/10.1038/s41531-020-00122-1.

(34) de Giorgi, F.,, Laferrière, F.,, Zinghirino, F.,, Faggiani, E.,, Lends, A.,, Bertoni, M.,, Yu, X.,, Grélard, A.,, Morvan, E.,, Habenstein, B.,, Dutheil, N.,, Doudnikoff, E.,, Daniel, J.,, Claverol, S.,, Qin, C.,, Loquet, A.,, Bezard, E.,, Ichas, F. Novel Self-Replicating α-Synuclein Polymorphs That Escape ThT Monitoring Can Spontaneously Emerge and Acutely Spread in Neurons. Sci. Adv. 2020, 6, eabc4364.

(35) Gath, J.,, Bousset, L.,, Habenstein, B.,, Melki, R.,, Böckmann, A.,, Meier, B. H. Unlike Twins: An NMR Comparison of Two α-Synuclein Polymorphs Featuring Different Toxicity. PLOS ONE 2014, 9 (3), e90659. https://doi.org/10.1371/JOURNAL.PONE.0090659.

(36) Landureau, M.,, Redeker, V.,, Bellande, T.,, Eyquem, S.,, Melki, R. The Differential Solvent Exposure of N-Terminal Residues Provides “Fingerprints” of Alpha-Synuclein Fibrillar Polymorphs. J. Biol. Chem. 2021, 296. https://doi.org/10.1016/J.JBC.2021.100737.

(37) Berezin, M. Y.,, Achilefu, S. Fluorescence Lifetime Measurements and Biological Imaging. Chem. Rev. 2010, 110 (5), 2641–2684. https://doi.org/10.1021/CR900343Z.

(38) Kafle, B. P. Molecular Luminescence Spectroscopy. Chem. Anal. Mat. Charac. Spectrophotometry 2020, 269–296. https://doi.org/10.1016/B978-0-12-814866-2.00009-9.

(39) Peng, C.,, Gathagan, R. J.,, Covell, D. J.,, Medellin, C.,, Stieber, A.,, Robinson, J. L.,, Zhang, B.,, Pitkin, R. M.,, Olufemi, M. F.,, Luk, K. C.,, Trojanowski, J. Q.,, Lee, V. M. Y. Cellular Milieu Imparts Distinct Pathological α-Synuclein Strains in α-Synucleinopathies. Nature 2018 557:7706 2018, 557 (7706), 558–563. https://doi.org/10.1038/s41586-018-0104-4.

(40) Guerrero-Ferreira, R.,, Taylor, N. M.,, Arteni, A.-A.,, Kumari, P.,, Mona, D.,, Ringler, P.,, Britschgi, M.,, Lauer, M. E.,, Makky, A.,, Verasdonck, J.,, Riek, R.,, Melki, R.,, Meier, B. H.,, Böckmann, A.,, Bousset, L.,, Stahlberg, H. Two New Polymorphic Structures of Human Full-Length Alpha-Synuclein Fibrils Solved by Cryo-Electron Microscopy. eLife 2019, 8, e48907. https://doi.org/10.7554/eLife.48907.

(41) Lautenschläger, J.,, Stephens, A. D.,, Fusco, G.,, Ströhl, F.,, Curry, N.,, Zacharopoulou, M.,, Michel, C. H.,, Laine, R.,, Nespovitaya, N.,, Fantham, M.,, Pinotsi, D.,, Zago, W.,, Fraser, P.,, Tandon, A.,, St George-Hyslop, P.,, Rees, E.,, Phillips, J. J.,, de Simone, A.,, Kaminski, C. F.,, Schierle, G. S. K. C-Terminal Calcium Binding of α-Synuclein Modulates Synaptic Vesicle Interaction. Nat. Commun. 2018, 9 (1), 1–13. https://doi.org/10.1038/s41467-018-03111-4.

(42) Han, J. Y.,, Choi, T. S.,, Kim, H. I. Molecular Role of Ca2+ and Hard Divalent Metal Cations on Accelerated Fibrillation and Interfibrillar Aggregation of α-Synuclein. Sci. Rep. 2018, 8 (1), 1–11. https://doi.org/10.1038/s41598-018-20320-5.

(43) Lövestam, S.,, Schweighauser, M.,, Matsubara, T.,, Murayama, S.,, Tomita, T.,, Ando, T.,, Hasegawa, K.,, Yoshida, M.,, Tarutani, A.,, Hasegawa, M.,, Goedert, M.,, Scheres, S. H. W. Seeded Assembly in Vitro Does Not Replicate the Structures of α-Synuclein Filaments from Multiple System Atrophy. FEBS Open Bio 2021, 11 (4), 999–1013. https://doi.org/10.1002/2211-5463.13110.

(44) Chen, S. W.,, Drakulic, S.,, Deas, E.,, Ouberai, M.,, Aprile, F. A.,, Arranz, R.,, Ness, S.,, Roodveldt, C.,, Guilliams, T.,, De-Genst, E. J.,, Klenerman, D.,, Wood, N. W.,, Knowles, T. P. J.,, Alfonso, C.,, Rivas, G.,, Abramov, A. Y.,, Valpuesta, J. M.,, Dobson, C. M.,, Cremades, N. Structural Characterization of Toxic Oligomers That Are Kinetically Trapped during α-Synuclein Fibril Formation. Proc. Natl. Acad. Sci. U.S.A. 2015, 112 (16), E1994–E2003. https://doi.org/10.1073/PNAS.1421204112.

