## Supporting Information for "Label-free characterisation of amyloids and alpha-Synuclein polymorphs by exploiting their intrinsic fluorescence property"

This PDF includes:

### Methods and Materials

#### Purification of αS

Human wild-type (WT) alpha-synuclein (αS) was expressed from plasmid pT7-7. The plasmid was heat shocked into Escherichia coli One Shot® BL21 STAR™ (DE3) (Invitrogen, Thermo Fisher Scientific, Cheshire, UK) and purified as previously described by periplasmic lysis of *E. coli* and chromatographic separation of the proteins.[1] Recombinant αS was purified using ion exchange chromatography (IEX) in buffer A (10 mM Tris, 1 mM EDTA pH 8) against a linear gradient of buffer B (10 mM Tris, 1 mM EDTA, 0.15 M $(NH_4)_2SO_4$ pH 8) on a HiPrep Q FF 16/10 anion exchange column (GE Healthcare, Uppsala, Sweden). The concentration of $(NH_4)_2SO_4$ in the pooled αS fractions was calculated and the equivalent $(NH_4)_2SO_4$ added to make the protein solution to 1 M. αS was then further purified on a HiPrep Phenyl FF 16/10 (High Sub) hydrophobic interaction chromatography (HIC) column (GE Healthcare) in buffer C (1 M $(NH_4)_2SO_4$, 50 mM Bis-Tris pH 7) and eluted against buffer D (50 mM Bis-Tris pH 7). Purification was performed on an ÄKTA Pure (GE Healthcare). αS was concentrated using 10 k MWCO amicon centrifugal filtration devices (Merck KGaA, Darmstadt, Germany) and monomeric αS isolated using a gel filtration HiLoad 16/60 superdex 75 pg column (GE Healthcare) in 20 mM Tris pH 7.4 and stored at −80 °C until use. Protein concentration was determined from the absorbance measurement at 280 nm on a Nanovue spectrometer using the extinction coefficient for αS of 5960 $M^{-1}cm^{-1}$.

#### Purification of TasA

*Bacillus subtilis* TasA was expressed using a pET24 plasmid. Residues 1-27, encoding the signal peptide of TasA was not included to permit purification of the mature $TasA^{28\text{-}261}$ protein, subsequently referred to as TasA. A 6xHis-tag was situated at the N terminus and a tobacco etch virus (TEV) protease recognition site ENLYFQ/x inserted before the N-terminus of the TasA protein to give the sequence MHHHHHHENLYFQ/F where '/' denotes the TEV cleavage site and F the beginning residue of the mature TasA. pET24:*Δ27tasA* was transformed into *E. coli* BL21 (DE3) pLysS cells which were grown at 37°C with constant shaking at 250 rpm to an optical density at 600 nm of 0.8. Protein expression was induced with 1 mM isopropyl-β-thiogalactopyranoside (IPTG) and grown overnight at 21°C with shaking at 200 rpm. *E. coli* cultures were centrifuged at 8000 rpm for 15 min and the pellet resuspended in 20 mM Tris, pH 8, 0.5 M NaCl, 1 tablet protease inhibitors, 1 mM $MgCl_2$ and sonicated 5 × 30 s on ice. The sonicate was centrifuged at 10,000 rpm, 4°C for 15 min. The supernatant was filtered through a 0.22 um filter and loaded onto a HisTrap Excel 5 mL column (GE Healthcare). The column was equilibrated in buffer A (20 mM Tris, 0.5 M NaCl, 30 mM imidazole, pH 8), and proteins were eluted through the column with a linear gradient to 100% of buffer B (20 mM Tris, 0.5 M NaCl, 0.5 M imidazole, pH 8). The purified His-tagged TasA protein was buffer exchanged using a PD-10 desalting column (GE Healthcare) equilibrated in TEV buffer (50 mM Tris-HCl (pH 8.0), 0.5 mM EDTA), followed by incubating with TEV protease (1 TEV: 50 TasA protein) at 30°C for 3 hours to cleave the His-tag. The TEV cleaved TasA was then eluted through the two stacked HisTrap Excel 5 mL columns in 20 mM Tris, 50 mM NaCl, pH 8, where the His-tag and TEV protein (containing its own His-tag) were retained on the column and the cleaved TasA eluted in the flow through. Protein concentration was determined from the absorbance measurement at 280 nm on a Nanovue spectrometer using the extinction coefficient for TasA of 14440 $M^{-1}$ $cm^{-1}$. The mass of purified proteins was analysed using electrospray ionisation mass spectrometry (ESI-MS) at the Department of Chemistry, University of Cambridge.

**Protein purity analysis by reversed phase chromatography**

RP chromatography was carried out on a 1260 Infinity high-pressure liquid chromatography (HPLC) system (Agilent Technologies LDA UK Ltd.) to measure the purity of αS and TasA. 50 μL of sample was injected onto a Discovery BIOWide Pore C18 column (15 cm × 4.6 mm, 5 μm column with a guard column) (Supelco, Merck) and eluted with a gradient of 95% water and 0.1% acetic acid and 5% acetonitrile and 0.1% acetic acid to 5% water and 0.1% acetic acid and 95% acetonitrile and 0.1% acetic acid at a flow rate of 0.8 mL $min^{-1}$ over 40 min. The elution profile was monitored by UV absorption at 220 and 280 nm. The area under the peaks in the chromatograph of absorption at 280 nm was calculated to provide the percentage purity. αS was 98% pure and TasA was 97% pure.

**Protein aggregation**

βLG (CAS 9045-23-2, Merck KGaA) powder was monomerised in hexa-iso-propanol (HFIP, Merck KGaA), before overnight lyophilisation (LyoQuest 85, Telstar, Spain). The freeze-dried monomers were dissolved in milli-Q water (MerckBurlington, MA, USA) to 100 μM. The mixture was then vortexed briefly and filtered through a 0.22 μm syringe filter (Millex-GS, Merck KGaA) to remove any clumps, before adjustment to pH2 using HCl (Merck, KGaA). Following this, the protein was aggregated at 80 °C for 3 hours on a ThermoMixer (Eppendorf, Hamburg, Germany). For storage, 0.05% of $NaN_3$ was added after βLG were formed. TasA was incubated at 70 μM at RT in 10 mM Tris pH 8 with 0.05% $NaN_3$ to prevent bacterial growth, rotating at 20 rpm (SB1, Stuart Scientific) for 1 week. αS was incubated at 100 μM in 10 mM Tris, or 140 mM KCl, 10 mM Tris pH 7.4 (mimicking intracellular conditions) or 140 mM NaCl, 1 mM CaCl2, 10 mM Tris (mimicking extracellular conditions) with 0.05% $NaN_3$ for 2 weeks at 37 °C rotating at 20 rpm on a (SB2, Stuart Scientific).

#### Circular dichroism (CD)

Protein samples were diluted to 2.5 μM and analysed in a 1mm cuvette at a temperature of 20°C. CD spectra were acquired using a JASCO J-810 spectropolarimeter (Jasco Inc, Easton, MD, USA). Spectra were recorded over the spectral range of 250 − 200 nm, with a resolution of 0.5 nm, a continuous scan at 50nm $min^{-1}$, and a bandwidth resolution of 1 nm. 10 accumulations were obtained for each sample and three preparations of each protein and buffer condition were measured. CD spectra of buffer only were recorded and subtracted from each sample spectrum. Mean residue ellipticity was calculated using **Equation 1**:

*Equation 1*

$$[\theta] = \frac{\theta_{obs}}{lcn}$$

where $[\theta]$ is the mean residue ellipticity (◦ $cm^2$ $dmol^{-1}$), $\theta_{obs}$ the observed ellipticity, $l$ the path length (mm), $c$ the molar concentration and $n$ the number of residues (i.e., 140 amino acids (a.a.) for αS, 233 a.a. for TasA and 162 a.a. for βLG).

#### Fluorescence characterisation

Protein samples were loaded into a cuvette at 100 μM at room temperature, and placed into a spectrophotometer (F-4500, Hitachi, Tokyo, Japan). Excitation and emission spectra were captured by emission at 440 nm over a frequency sweep between 280—420 nm (at 1 nm intervals), and excitation at 370 nm over 400—500 nm (at 1 nm intervals), respectively. Excitation and emission slits of 10 mm, and a scan speed of 240 nm $min^{-1}$ were used. The fluorescence spectra of the buffer were measured at the same settings as the protein and subtracted from the final spectra. Measurements were based on triplicate measurements of three individual protein preparations. For the final plot, a MATLAB (MathWorks, Natick, MA, USA) script was used to identify the peak excitation and emission wavelengths and normalise the spectra across a range of 16 nm centred on the peak values.

**Two-photon (2P-) fluorescence lifetime imaging microscopy (FLIM)**

Samples at 100 µM were centrifuged and washed in $dH_2O$ three times to remove salts. For HFIP treatment, 20 µL of HFIP (Merck KGaA) was added to 10 µL of 100 µM protein solution. After pipetting the sample to mix, the mixture was dried with a stream of $N_2$, before resuspension in 10 µL $dH_2O$. Protein samples were deposited onto a 1.5 thickness precision coverslip (Marienfeld GmbH, Lauda-Konigshofen, Germany) and dried on a hotplate for ~10 mins.

Samples were imaged on a home-built confocal fluorescence microscope equipped with a time-correlated single photon counting (TCSPC) module. A pulsed, femtosecond Ti:S laser (MaiTai DeepSee, SpectraPhysics, Oxford, UK) provided excitation at 740 nm and a repetition rate of 80 MHz. This was passed into a commercial microscope frame (IX83, Olympus, Tokyo, Japan) through a 60x oil objective (PlanApo 60XOSC2, 1.4 NA, Olympus). A bandpass filter of 450/50 (Chroma Technologies, Rockingham, VT, USA) was applied to the 2P emission to separate it from the excitation light. Laser scanning was performed using a galvanometric mirror system (Quadscanner, Aberrior, Gottingen, Germany). Emission photons were collected on a photon multiplier tube (PMT, PMC150, B&H GmBH, Berlin, Germany) and relayed to a time-corelated single photon counting card (SPC830, B&H GmBH). Images were acquired at 256x256 pixels for 200 s (i.e., 20 cycles of 10 s). Photon counts were kept below 1% of SYNC rates to prevent photon pile-up. TCSPC images were analysed using an in-house phasor plot analysis script (https://github.com/LAG-MNG-CambridgeUniversity/TCSPCPhasor), from which lifetime maps and phasor plots were generated. A general introduction to phasor plots is given in **Supplementary Page 9**.

**Atomic force microscopy (AFM)**

For imaging individual fibrils for morphological analysis, protein solutions were diluted to 10 μM in $dH_2O$, and incubated on poly-l-lysine (Merck KGaA) coated mica for 30 mins. To remove salts, the mica was washed thrice with $dH_2O$ and dried under a gentle stream of $N_2$. The protein "meshes" corresponding to those imaged using 2P were prepared as described above. Imaging was performed on a BioScope Resolve (Bruker GmbH, Karlsruhe, Germany). The instrument was operated in ScanAsyst Air mode with a silicon nitride tip of a spring constant 40 $N\,m^{-1}$ and nominal tip diameter of 2 nm (SCANASYST-AIR, Bruker). Images were collected at a scan rate of 1 Hz and resolution of 512x512 pixels.

Acquired AFM images were computationally flattened on Nanoscope Analysis 9.4 (Bruker GmBH) before import into MATLAB (MathWorks) using the MATLAB toolkit for Bruker. Batch analysis was performed using an in-house MATLAB script. A combination of manual and automatic segmentation of individual fibrils was performed. The height profile from each fibril was smoothed and characterised as either smooth or periodic as by a peak/trough search algorithm. For calculation of average fibril height, the mean across the whole fibril length and of the peaks were used for smooth and periodic fibrils respectively. For periodic fibrils, the mean height of its peaks and troughs were additionally calculated.

**Limited proteolysis of αS fibrils**

100 μM of αS fibrils were incubated at 37°C in 3.8 $\mu g\,mL^{-1}$ Proteinase K. 10 μL aliquots were removed at time points, 0, 1, 5 and 15 minutes and incubated with 20 mM PMSF to inactivate the proteinase K. The samples were frozen and lyophilised using a LyoQuest 85 freeze-dryer (Telstar, Spain). The protein films were solubilised in HFIP. HFIP was then evaporated under a stream of $N_2$ and the samples resuspended in LDS buffer before being

heated to 100°C and analysed by SDS-PAGE on a 4-12% Bis-Tris gel (NuPAGE, Thermo Scientific) and stained with Coomassie blue (Merck KGaA).

**Plotting and statistical analysis**

All statistical analyses were performed on Prism 6 (GraphPad, San Diego, CA, USA), where a one-way ANOVA test with Holm-Sidak's multiple comparison was applied. Violin plots were produced using MATLAB (MathWorks) using open-source code from Anne Urai (github.com/anne-urai).

### Supplementary figures

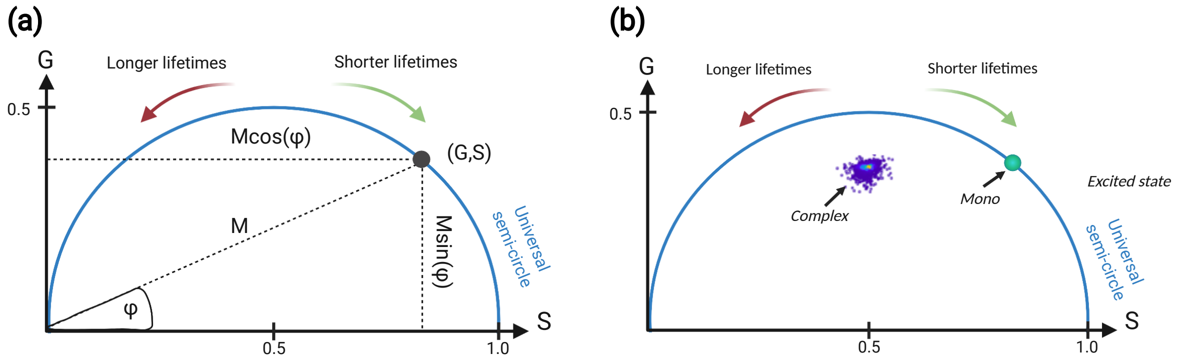

**Supplementary Figure 1: Representative phasor plot.** (a) Time-domain TCSPC data can be Fourier-transformed into the frequency-domain (FD) and represented as 'phasors' with (G,S) coordinates on a polar plot (i.e., phasor plot). Analogous to FD-FLIM, resulting phasors have an associated phase (φ) and modulation (M), shifts in which indicate changes in fluorescence lifetimes. (b) Mono-exponential and complex exponential phasors fall on and within the universal semicircle respectively. Created on BioRender.com.

Phasor plots are an efficient way to represent complex exponential decay fluorescence lifetimes.[2–4] It is a model-free, global representation of fluorescence lifetime data, which clearly shows how ensemble image pixels are affected by the chemical, physical and environmental changes of the sample. It stems from frequency-domain (FD-)FLIM where an intensity modulated light source is used to excite the sample, and the demodulation and phase shift in the resulting fluorescence emission is measured. Hence, the phase angle (φ) and modulation (M) for each pixel within the image can be conveniently represented on a phasor plot, with real and complex coordinates, (G,S).[2–5] For use with time-domain time-correlated single photon counting (TCSPC) data, a Fourier transform is required. The exponential decays of each of the 256 time bins, along each 256 by 256 x- and y-coordinates of the TCSPC image is represented as a single plot (i.e., 'phasor') within the plot (**Supplementary Figure 1a**), with (G,S) calculated as follows:

*Equation 2*

$$G(\omega) = \frac{\int_0^\infty I(t)\cos(\omega t)\,dt}{\int_0^\infty I(t)dt}$$

*Equation 3*

$$S(\omega) = \frac{\int_0^\infty I(t)\sin(\omega t)\,dt}{\int_0^\infty I(t)dt}$$

where $\omega = 2\pi f$ is the laser repetition angular frequency, $f$ is the laser repetition rate, $I$ is fluorescence intensity (i.e. total number of photons) and $t$ is time. The phasor plot has a universal semicircle, which represents mono-exponential decay lifetimes (**Equation 4, Supplementary Figure 1b**):

*Equation 4*

$$(G(\omega) - 0.5)^2 + S(\omega)^2 = 0.25$$

This is dependent on $\omega$, hence scales with $f$. Calibration of the phasor plot was performed using standard fluorescence dyes of known mono-exponential fluorescence lifetimes, i.e., Coumarin 6 (Merck KGaA) in ethanol. Multi-exponential species (e.g., intrinsic amyloid fluorescence) have phasors lying within the universal semicircle (**Supplementary Figure 1b**). An independent determination of fluorescence lifetime as modulation ($\tau_M$, **Equation 5**) and phase ($\tau_\phi$, **Equation 6**) can be made:

*Equation 5*

$$\tau_\phi = \frac{S(\omega)}{\omega\, G(\omega)}$$

*Equation 6*

$$\tau_M = \frac{1}{\omega}\sqrt{\frac{\cos(\phi)^2}{G(\omega)^2} - 1} = \frac{1}{\omega}\sqrt{\frac{\sin(\phi)^2}{S(\omega)^2} - 1}$$

where $\phi = \tan^{-1}\left(\frac{S}{G}\right)$ is the phase. For purposes of comparison in the manuscript, $\tau_M$ (i.e., more sensitive to changes in G) is used over $\tau_\phi$ (i.e., more sensitive to changes in S) as it is the more sensitive fluorescence lifetime for changes between different amyloid/polymorph samples.

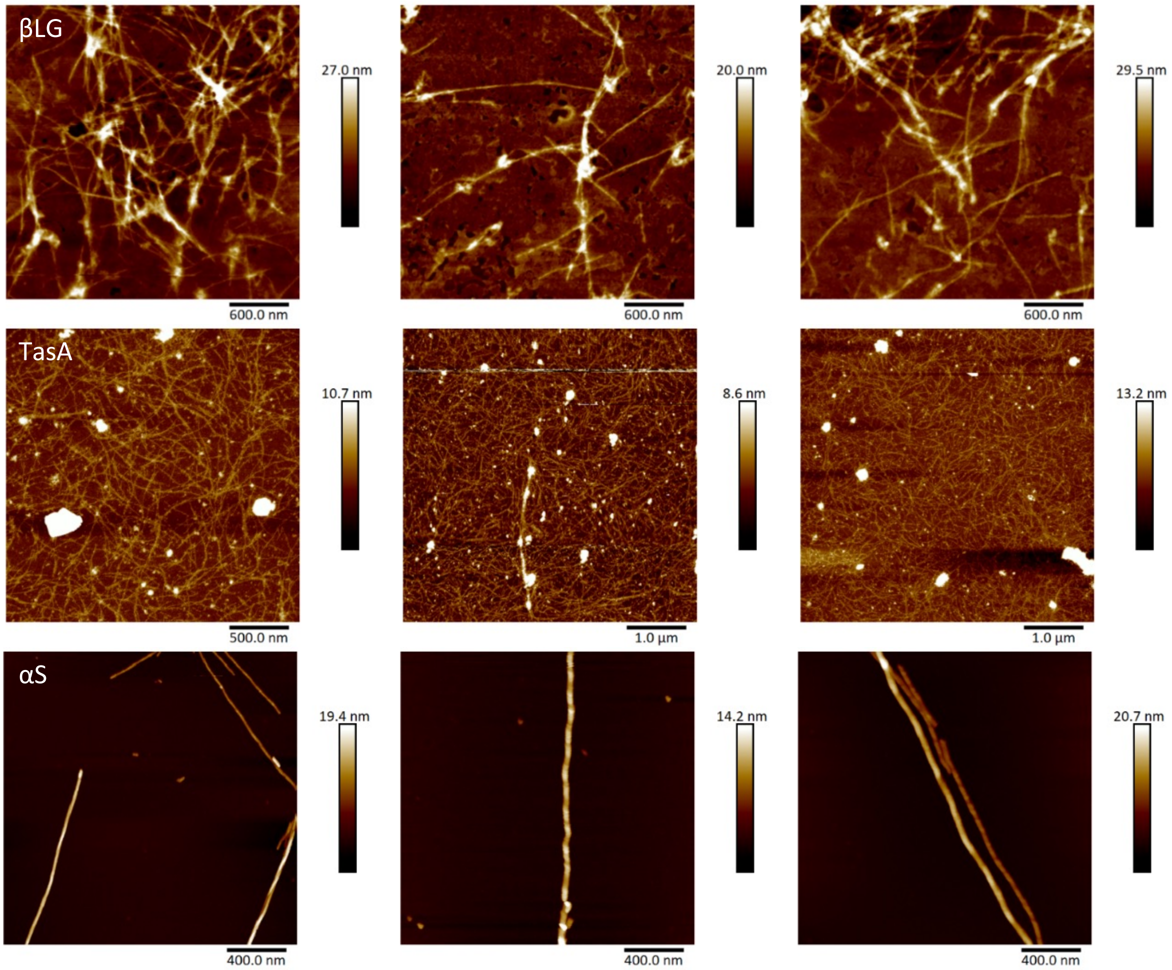

**Supplementary Figure 2: βLG, TasA and αS feature different fibril morphologies.** Representative AFM images of 10 µM of each protein. βLG and TasA have no/less periodicity compared to αS fibrils. TasA fibrils are very thin in comparison to βLG and αS. Height profile is shown in brown scale.

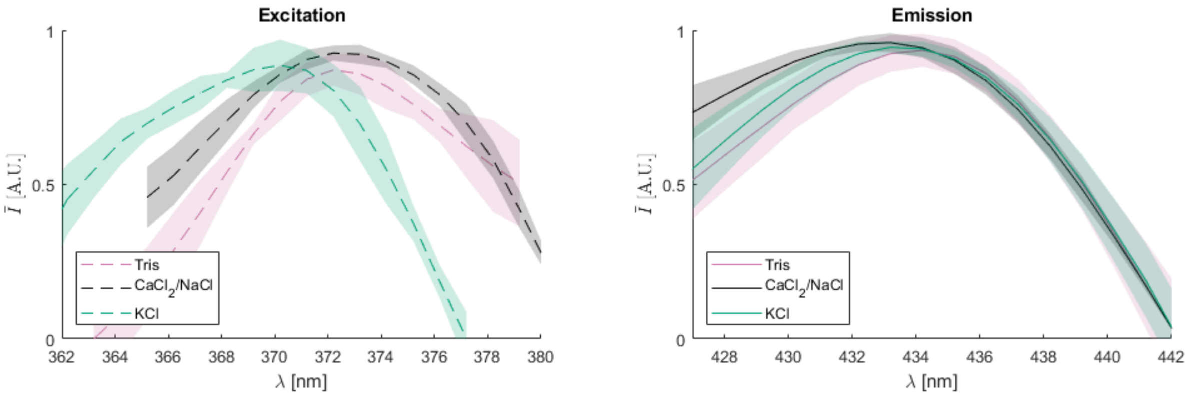

**Supplementary Figure 3: αS aggregated in different salt buffers have similar excitation and emission profiles.** (a) Excitation and emission spectra were measured between 340 to 400 nm with the emission set at peak emission, and emission spectra were measured between 400 to 460 nm with the excitation set at peak excitation for each protein. Excitation and emission peaks for each protein were: Tris – ex 373 nm, em 433 nm, $CaCl_2$/NaCl – ex 373 nm, em 433 nm, and KCl – 370 nm, em 432 nm. Displayed are the average of three scans from three individual protein preparations.

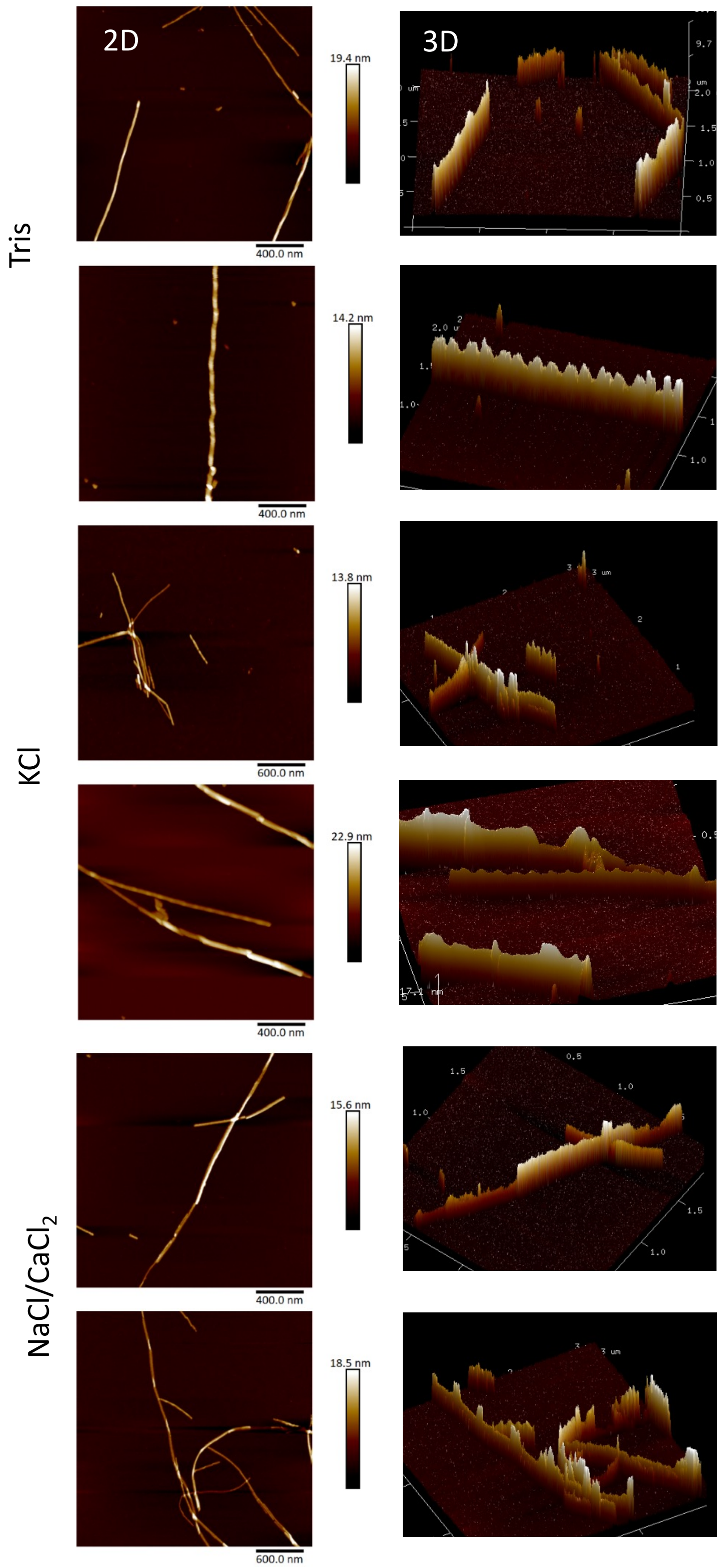

**Supplementary Figure 4: Different fibril polymorphs are observed for aSyn in different buffers.** Representative AFM images of 5 µM αS in 20 mM Tris pH 7.4, with additional KCl (140 mM) or NaCl (140 mM) and $CaCl_2$ (1 mM) on freshly cleaved mica. 2D images on the left show the height profile of fibrils. The 3D images on the right more clearly show periodicity of the fibrils.

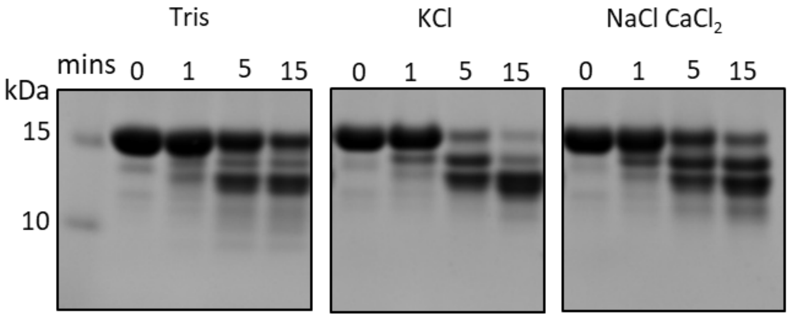

**Supplementary Figure 5. Repeated limited proteolysis profiles of αS fibrils shows reproducibility in digestion profiles.** 100 µM of αS in each buffer condition was incubated in 3.8 µg/ml proteinase K for 0, 1, 5 and 15 mins. Monomeric αS is ~14.4 kDa, degradation patterns of the three samples show a similar band profile, but the intensities differ, indicating differences in cleavage rates. This repeat has similar proteolysis profiles to **Figure 4**, showing reproducible results.

### References

(1) Stephens, A. D.; Matak-Vinkovic, D.; Fernandez-Villegas, A.; Kaminski Schierle, G. S. Purification of Recombinant α-Synuclein: A Comparison of Commonly Used Protocols. *Biochem.* **2020**, *59* (48), 4563–4572. https://doi.org/10.1021/ACS.BIOCHEM.0C00725

(2) Digman, M. A.; Caiolfa, V. R.; Zamai, M.; Gratton, E. The Phasor Approach to Fluorescence Lifetime Imaging Analysis. *Biophys. J.* **2008**, *94* (2), L14. https://doi.org/10.1529/BIOPHYSJ.107.120154.

(3) Schlachter, S.; Elder, A. D.; Esposito, A.; Kaminski, G. S.; Frank, J. H.; van Geest, L. K.; Kaminski, C. F.; Matthews, S. M.; Yunus, K.; Brennan, C. M.; Fisher, A. C. MhFLIM: Resolution of Heterogeneous Fluorescence Decays in Widefield Lifetime Microscopy. *Opt. Express* **2009**, *17* (3), 1557–1570. https://doi.org/10.1364/OE.17.001557.

(4) Ranjit, S.; Malacrida, L.; Jameson, D. M.; Gratton, E. Fit-Free Analysis of Fluorescence Lifetime Imaging Data Using the Phasor Approach. *Nat. Protoc.* **2018**, *13* (9), 1979–2004. https://doi.org/10.1038/s41596-018-0026-5.

(5) Elder, A. D.; Kaminski, C. F.; Frank, J. H. Φ2FLIM: A Technique for Alias-Free Frequency Domain Fluorescence Lifetime Imaging. *Opt. Express* **2009**, *17* (25), 23181–23203. https://doi.org/10.1364/OE.17.023181.
